## Supplementary material for "The dorsal fan-shaped body is a neurochemically heterogeneous sleep-regulating center in *Drosophila*": S1 Table

| **FBS line** | **AD component** | **Average whole brain dFB neurons ± SEM (n)** | **Range dFB neurons** | **Bowtie VNC-SP (Y/N)** | **Range of additional cells in brain** | **Average VNC cells in metathoracic ganglion ± SEM (n)** | **TPN1 (Y/N)** | **Range of additional cells in VNC** |
| --- | --- | --- | --- | --- | --- | --- | --- | --- |
| FBS 1 | 12D12-AD | 4.64 ± 0.40 (n=14) | 2-7 | Y | 0 | 2.00 ± 0.32 (n=11) | N | 0 |
| FBS 6 | 42H01-AD | 3.85 ± 0.52 (n=13) | 1-4 | N | 0 | 0.00 ± 0.00 (n=8) | N | 0 |
| FBS 25 | 22A12-AD | 0 ± 0 (n=5) | 0 | N | 0 | 0.00 ± 0.00 (n=5) | N | 0 |
| FBS 28 | 54G07-AD | 13.08 ± 0.63 (n=12) | 10-18 | Y | 0-1 | 3.09 ± 0.17 (n=22) | N | 0 |
| FBS 33 | 49C04-AD | 26.50 ± 1.15 (n=10) | 23-31 | Y | 0 | 3.17 ± 0.32 (n=12) | Y | 0 |
| FBS 35 | 65C03-AD | 5.00 ± 1.06 (n=6) | 2-8 | Y | 1-2 | 3.60 ± 0.24 (n=5) | Y | 0 |
| FBS 41 | VT058545-AD | 11.50 ± 0.43 (n=10) | 10-14 | N | 0-2 | 3.00 ± 0.41 (n=4) | Y | 0 |
| FBS 42 | 12G09-AD | 21.56 ± 0.80 (=10) | 18-25 | Y | 2-4 | 3.00 ± 0.37 (n=6) | Y | 0-2 |
| FBS 45 | VT050238-AD | 25.11 ± 1.41 (n=9) | 21-32 | Y | 0 | 2.40 ± 0.16 (n=15) | N | 0 |
| FBS 53 | VT043400-AD | 23.86 ± 0.51 (n=7) | 22-26 | Y | 0 | 2.91 ± 0.31 (n=11) | N | 0-1 |
| FBS 57 | VT025720-AD | 7.40 ± 0.68 (n=5) | 5-9 | Y | 0 | 3.00 ± 0.31 (n=7) | Y | 0 |
| FBS 58 | VT064566-AD | 21.50 ± 1.28 (n=6) | 18-27 | Y | 0-2 | 3.10 ± 0.28 (n=10) | Y | 0-1 |
| FBS 60 | VT023818-AD | 24.56 ± 0.50 (n=9) | 22-27 | Y | 0 | 3.27 ± 0.38 (n=11) | Y | 0-2 |
| FBS 64 | VT027956-AD | 15.00 ± 2.12 (n=7) | 11-19 | Y | 0 | 3.40 ± 0.49 (n=10) | Y | 0 |
| FBS 68 | VT023823-AD | 4.45 ± 0.34 (n=11) | 3-6 | Y | 0 | 4.08 ± 0.49 (n=13) | Y | 0-4 |
| FBS 70 | VT034811-AD | 4.86 ± 0.40 (n=7) | 3-6 | Y | 0-7 | 3.90 ± 0.38 (n=10) | Y | 0-4 |
| FBS 72 | VT026953-AD | 24.40 ± 1.50 (n=7) | 20-28 | Y | 0-1 | 3.27 ± 0.25 (n= 10) | Y | 0 |
| FBS 81 | 74G10-AD | 22.17 ± 0.70 (n=6) | 19-24 | Y | 0 | 2.80 ± 0.58 (n=5) | N | 0-2 |
| FBS 84 | VT007754-AD | 23.80 ± 0.58 (n=8) | 22-25 | Y | 0-2 | 3.36 ± 0.31 (n=12) | Y | 0-4 |
| FBS 87 | VT045793-AD | 18.55 ± 0.53 (n=13) | 17-22 | Y | 0-2 | 3.62 ± 0.29 (n=13) | Y | 0-6 |
| dFB-Split | 84C10-AD | 23.11 ± 0.68 (n=18) | 18-27 | N | 0 | 0.00 ± 0.00 (n=18) | N | 0 |
