## Supplementary material for "The dorsal fan-shaped body is a neurochemically heterogeneous sleep-regulating center in *Drosophila*": S2 Table

|  | **Total Sleep Change-Thermogenetic** | | **Daytime Bout Duration-Thermogenetic** | | **Nighttime Bout Duration-Thermogenetic** | | **Total Sleep Change-Optogenetic** | | **Daytime Bout Duration-Optogenetic** | | **Nighttime Bout Duration-Optogenetic** | |
| --- | --- | --- | --- | --- | --- | --- | --- | --- | --- | --- | --- | --- |
| **FBS line** | **Females** | **Males** | **Females** | **Males** | **Females** | **Males** | **Females** | **Males** | **Females** | **Males** | **Females** | **Males** |
| **FBS 1** |  |  |  |  |  |  |  |  |  |  |  |  |
| **FBS 6** |  |  |  |  |  |  |  |  |  |  |  |  |
| **FBS 25** |  |  |  |  |  |  |  |  |  |  |  |  |
| **FBS 28** |  |  | **X** |  |  |  |  |  |  |  |  |  |
| **FBS 33** |  |  |  |  |  |  |  | **X** |  | **X** |  |  |
| **FBS 35** |  |  |  |  |  |  |  |  |  | **X** |  |  |
| **FBS 41** |  |  |  | **(-)** |  |  |  |  |  |  |  |  |
| **FBS 42** | **X** | **X** | **X** | **X** |  | **(+)** | **X** | **X** | **X** | **X** | **X** | **X** |
| **FBS 45** | **X** | **X** | **X** | **X** | **(+)** | **X** | **X** | **X** | **X** | **X** | **X** | **X** |
| **FBS 53** | **X** | **X** | **X** | **X** | **(+)** |  | **X** | **X** | **X** | **X** | **X** | **X** |
| **FBS 57** |  |  |  |  |  |  |  |  | **X** |  |  |  |
| **FBS 58** | **(-)** |  |  |  |  |  |  |  |  |  | **X** |  |
| **FBS 60** |  |  |  |  |  |  |  |  | **X** |  |  |  |
| **FBS 64** | **(-)** |  |  |  |  |  |  |  |  | **X** |  |  |
| **FBS 68** | **X** | **X** | **X** | **X** | **(+)** | **X** | **X** | **X** | **X** | **X** | **X** | **X** |
| **FBS 70** |  |  |  |  |  |  | **X** |  |  |  |  |  |
| **FBS 72** |  | **X** | **X** | **X** |  |  | **X** | **X** | **X** | **X** | **X** | **X** |
| **FBS 81** |  | **X** |  |  |  | **(+)** |  | **X** | **X** | **X** |  | **X** |
| **FBS 84** |  | **X** | **X** | **X** |  |  |  | **X** | **X** | **X** |  |  |
| **FBS 87** |  |  |  |  |  |  |  |  |  | **X** |  |  |
