## Supplementary material for "The dorsal fan-shaped body is a neurochemically heterogeneous sleep-regulating center in *Drosophila*": S3 Table

| Genotype | Source or reference | Identifiers | Additional information |
| --- | --- | --- | --- |
| P{GMR23E10-GAL4}attP2 | Bloomington Drosophila Stock Center | RRID:BDSC_49032 |  |
| w[*]; P{y[+t7.7] w[+mC]=UAS-TrpA1(B).K}attP16 | Bloomington Drosophila Stock Center | RRID:BDSC_26263 | UAS-TrpA1 |
| Canton-S | Laboratory of Paul Shaw |  |  |
| w[*]; P{y[+t7.7] w[+mC]=10XUAS-IVS-mCD8::GFP}attP40 | Bloomington Drosophila Stock Center | RRID:BDSC_32186 |  |
| w[*]; P{y[+t7.7] w[+mC]=10XUAS-IVS-mCD8::GFP}attP2 | Bloomington Drosophila Stock Center | RRID:BDSC_32185 |  |
| w[1118]; P{y[+t7.7] w[+mC]=23E10-GAL4.DBD}attP2/TM3 | Bloomington Drosophila Stock Center | RRID:BDSC_69269 |  |
| w[1118]; P{y[+t7.7] w[+mC]=p65.AD.Uw}attP40 | Bloomington Drosophila Stock Center | RRID:BDSC_71210 | Empty-AD |
| w[1118]; P{y[+t7.7] w[+mC]=R84C10-p65.AD}attP40;MKRS/TM6B | Bloomington Drosophila Stock Center | RRID:BDSC_70833 | AD for dFB-Split |
| w[1118]; P{y[+t7.7] w[+mC]=20XUAS-IVS-CsChrimson.mVenus}attP40 | Bloomington Drosophila Stock Center | RRID:BDSC_55135 | Cs-Chrimson |
| w[1118]; P{y[+t7.7] w[+mC]=VT050238-p65.AD}attP40 | Bloomington Drosophila Stock Center | RRID:BDSC_72962 | AD for FBS45 |
| P{VT023823-p65.AD}attP40 | Bloomington Drosophila Stock Center | RRID:BDSC_86629 | AD for FBS68 |
| P{R12D12-p65.AD}attP40 | Bloomington Drosophila Stock Center | RRID:BDSC_70539 | AD for FBS1 |
| P{R42H01-p65.AD}attP40 | Bloomington Drosophila Stock Center | RRID:BDSC_70687 | AD for FBS6 |
| P{R22A12-p65.AD}attP40 | Bloomington Drosophila Stock Center | RRID:BDSC_70120 | AD for FBS25 |
| P{R54G07-p65.AD}attP40 | Bloomington Drosophila Stock Center | RRID:BDSC_70737 | AD for FBS28 |
| P{R49C04-p65.AD}attP40 | Bloomington Drosophila Stock Center | RRID:BDSC_71075 | AD for FBS33 |
| P{R65C03-p65.AD}attP40 | Bloomington Drosophila Stock Center | RRID:BDSC_71005 | AD for FBS35 |
| P{VT058545-p65.AD}attP40 | Bloomington Drosophila Stock Center | RRID:BDSC_74090 | AD for FBS41 |
| P{R12G09-p65.AD}attP40 | Bloomington Drosophila Stock Center | RRID:BDSC_68826 | AD for FBS42 |
| P{VT043400-p65.AD}attP40 | Bloomington Drosophila Stock Center | RRID:BDSC_75874 | AD for FBS53 |
| P{VT025720-p65.AD}attP40 | Bloomington Drosophila Stock Center | RRID:BDSC_73432 | AD for FBS57 |
| P{VT064566-p65.AD}attP40 | Bloomington Drosophila Stock Center | RRID:BDSC_71451 | AD for FBS58 |
| P{VT023818-p65.AD}attP40 | Bloomington Drosophila Stock Center | RRID:BDSC_73262 | AD for FBS60 |
| P{VT027956-p65.AD}attP40 | Bloomington Drosophila Stock Center | RRID:BDSC_73061 | AD for FBS64 |
| P{VT034811-p65.AD}attP40 | Bloomington Drosophila Stock Center | RRID:BDSC_71337 | AD for FBS70 |
| P{VT026953-p65.AD}attP40 | Bloomington Drosophila Stock Center | RRID:BDSC_74413 | AD for FBS72 |
| P{R74G10-p65.AD}attP40 | Bloomington Drosophila Stock Center | RRID:BDSC_71131 | AD for FBS81 |
| P{VT007754-p65.AD}attP40 | Bloomington Drosophila Stock Center | RRID:BDSC_74138 | AD for FBS84 |
| P{VT045793-p65.AD}attP40 | Bloomington Drosophila Stock Center | RRID:BDSC_71441 | AD for FBS87 |
| P{VT013602-p65.AD}attP40; P{23E10-GAL4.-DBD}attP2 | Jones et al. 2023 |  | VNC-SP Split |
| w[*]; P{y[+t7.7] w[+mC]=13XLexAop2-KZip+.3XHA}su(Hw)attP5/CyO; TM6B, Tb[1]/MKRS | Bloomington Drosophila Stock Center | RRID:BDSC_76253 |  |
| w[*]; P{y[+t7.7] w[+mC]=13XLexAop2-KZip+.3XHA}attP2 | Bloomington Drosophila Stock Center | RRID:BDSC_76254 |  |
| w[*]; P{w[+mC]=UAS-Hsap\KCNJ2.EGFP}7 | Bloomington Drosophila Stock Center | RRID:BDSC_6595 | UAS-Kir2.1 |
| pJFRC100-20XUAS-TTS-Shibire-ts1-p10 in VK00005 | Laboratory of Gerry Rubin |  | UAS-Shi^ts1^ |
| w[*]; Mi{Trojan-lexA:QFAD.0}ChAT[MI04508-TlexA:QFAD.0] CG7715[MI04508-TlexA:QFAD.0-X]/TM6B, Tb[1] | Bloomington Drosophila Stock Center | RRID:BDSC_60319 | ChAT-LexA |
| w[*]; TI{2A-p65(AD)::Zip+}VGlut[2A-p65AD]/Cyo | Bloomington Drosophila Stock Center | RRID:BDSC_84713 | VGlut-AD |
| w[*]; Mi{Trojan-p65AD.2}VGlut[MI04979-Tp65AD.2]/CyO | Bloomington Drosophila Stock Center | RRID:BDSC_82986 | VGlut-AD |
| w[*]; l(2)*/Cyo; Mi{Trojan-GAL4DBD.0}ChAT[MI04508-TG4DBD.0] CG7715[MI04508-TG4DBD.0-X]/TM3, Sb[1] | Bloomington Drosophila Stock Center | RRID:BDSC_60318 | ChAT-DBD |
| y[1] w[*]; Mi{Trojan-p65AD.2}Gad1[MI09277-Tp65AD.2]/TM6B, Tb[1] | Bloomington Drosophila Stock Center | RRID:BDSC_60322 | Gad1-AD |
| wg[Sp-1]/CyO; Mi{Trojan-GAL4DBD.2}Gad1[MI09277-TG4DBD.2]/TM6B, Tb[1] | Bloomington Drosophila Stock Center | RRID:BDSC_82987 | Gad1-DBD |
| w[1118]; P{y[+t7.7] w[+mC]=R84C10-GAL4.DBD}attP2 | Bloomington Drosophila Stock Center | RRID:BDSC_69553 | 84C10-DBD |
| w[*]; P{y[+t7.7] w[+mC]=10XUAS-EGFP.DD}attP2 | Bloomington Drosophila Stock Center | RRID:BDSC_79025 | GFP-DD |
| y[1] sc[*] v[1] sev[21]; P{y[+t7.7] v[+t1.8]=TRiP.HMS02175}attP40 | Bloomington Drosophila Stock Center | RRID:BDSC_40927 | VGlut RNAi |
| y[1] sc[*] v[1] sev[21]; P{y[+t7.7] v[+t1.8]=TRiP.HMS02011}attP2 | Bloomington Drosophila Stock Center | RRID:BDSC_40845 | VGlut RNAi |
| y[1] v[1]; P{y[+t7.7] v[+t1.8]=TRiP.JF02764}attP2 | Bloomington Drosophila Stock Center | RRID:BDSC_27684 | VAChT RNAi |
| y[1] v[1]; P{y[+t7.7] v[+t1.8]=TRiP.HMS06015}attP40 | Bloomington Drosophila Stock Center | RRID:BDSC_80435 | VAChT RNAi |
| y[1] v[1]; P{y[+t7.7] v[+t1.8]=UAS-GFP.VALIUM10}attP2 | Bloomington Drosophila Stock Center | RRID:BDSC_35786 | GFP RNAi |
| w[1118]; P{p65-AD.Uw}attP40; P{GAL4-DBD.Uw}attP2 | Bloomington Drosophila Stock Center | RRID:BDSC_79603 | Empty-Split |
| w[*]; P{y[+t7.7] w[+mC]=dj-GFP.S}AS1/Cyo | Bloomington Drosophila Stock Center | RRID:BDSC_5417 | Don Juan GFP |
